# Neuroanatomical signatures of depression and anxiety in at-risk adolescents: A symptom-oriented perspective

**DOI:** 10.1101/2025.11.23.689891

**Authors:** Qingwen Ding, Divyangana Rakesh, Siya Peng, Xinying Li, Sarah Whittle

## Abstract

**Background:** Major depressive disorder (MDD) and generalized anxiety disorder (GAD) often emerge in adolescence. Identifying their symptom structures and neural biomarkers in youth with elevated risk is critical for early detection and developing interventions. This study investigated the central symptoms of MDD and GAD and identified the key brain regions (“brain bridges”) linking structural neuroanatomy to symptoms in at-risk adolescents.

**Methods:** A total of 1,568 adolescents at high risk for MDD and 413 adolescents at high risk for GAD were identified from the Adolescent Brain Cognitive Development (ABCD) study. Participants completed MDD and GAD symptom assessments and underwent Magnetic Resonance Imaging scanning. We constructed 10 psychological network models incorporating symptoms and brain structural indices to identify central symptoms and brain bridges.

**Results:** Concentration difficulties emerged as the shared core symptom, while self-hatred and irritability were the unique central symptoms for MDD and GAD, respectively. Left- and right-pallidum volumes exhibited the highest bridge centrality for MDD symptoms, whereas left-nucleus accumbens volume showed the highest bridge centrality for GAD symptoms.

**Conclusion:** Our findings highlight the pallidum and nucleus accumbens as key brain regions linking structural anatomy to symptom networks in high-risk youth. These results provide novel insights into early neurobiological markers of MDD and GAD from a symptom-oriented perspective and may inform targeted interventions.

## Introduction

Major depressive disorder (MDD) and generalized anxiety disorder (GAD) are among the most common mental disorders and are highly comorbid (1). Both disorders are heterogeneous in their clinical presentation, encompassing a wide range of symptoms across affective, cognitive, neurovegetative, and psychomotor domains, and many of them overlap (e.g., concentration difficulties, fatigue, and sleep problems). The high comorbidity and shared symptomatology raise an important debate about whether MDD and GAD should be classified as distinct disorders (2). To address this debate, a closer examination of their symptom structure and neural underpinnings is needed. This investigation is particularly crucial in adolescence, as depression and anxiety symptoms frequently emerge during this critical developmental period (3,4). If left untreated, adolescents with elevated symptoms will face an increased risk of long-term poor mental health and functioning (5,6). Identifying the neurobiological mechanisms underlying these symptoms in a high-risk group may help uncover early biomarkers. This knowledge could also inform targeted preventive interventions designed to improve individual outcomes and reduce long-term healthcare costs.

A popular approach to explore the structure of psychiatric symptoms is the recently developed psychological network model (7). This model views disorders as arising from the dynamic interplay between individual symptoms (8). By conceptualizing symptoms within a psychiatric disorder as nodes and their associations as edges, the network model can reveal reciprocal dependencies between symptoms. Central symptoms, which share stronger relationships with other symptoms, are integral to the network structure and are thought to play a crucial role in the development and maintenance of the disorder (9). Targeting these central symptoms in interventions is thought to lead to a widespread reduction in symptom severity, thereby effectively reducing risk and alleviating the overall burden of the disorder. A substantial body of research in youth samples has identified the central symptoms in depression and anxiety (e.g., self-hatred and uncontrolled worries) (10–14). However, they have primarily focused on general populations. Identifying central symptoms in high-risk youth may specifically reveal key factors contributing to their increased vulnerability, offering prioritized targets for early intervention.

Extensive research has also explored the neural markers of youth depression and anxiety (15–23). However, findings have generally been mixed, with conflicting results reported (21–23). For instance, some studies examining the nucleus accumbens (NAcc), a key region involved in reward processing, have linked decreased volume to anxiety (17) and depression (20), while others have associated increased volume with depression (18) or found no significant associations (19). Studies examining anxiety and depression simultaneously have reported no shared neuroanatomical substrates, further complicating interpretations (15,19). One key reason for these discrepancies is the reliance on aggregate symptom scores, which treats symptoms as interchangeable in their associations with brain structure. Given the heterogeneity of MDD and GAD, individual symptoms may have distinct neurobiological correlates (24,25). Analyzing symptoms individually, rather than aggregating them, could provide a more nuanced understanding of the neurobiological correlates of these disorders (26). However, the neural markers of GAD and MDD at the symptom level still remain unknown.

A promising approach to addressing this complexity and heterogeneity is the integration of brain structures as nodes within psychiatric symptom networks, forming a “brain-symptom network.” This approach bridges brain anatomy and symptom expression, offering deeper and more comprehensive insights into the neural correlates of these disorders. First, the integrated network model enables the identification of intricate associations between individual symptoms under a psychiatric construct and regional brain structures. It allows for more precise estimation of brain-symptom edges by accounting for all other potential associations within the network. Second, the model calculates brain-symptom edges to derive accumulative bridge centrality indices. These indices can determine which brain regions play a key role in linking structural anatomy to psychiatric symptoms, referred to as “brain bridges.” Similar to central symptoms in symptom-only networks, brain bridges may play a central role in the development and maintenance of psychiatric networks and represent targets for intervention. Finally, by locating the specific symptoms to which these brain bridges are connected, this approach can delineate the primary pathways through which brain regions contribute to the disorder.

This study aimed to identify the symptom structures of MDD and GAD and their neuroanatomic signatures in adolescents at high risk (i.e., presenting with subthreshold symptoms) from the Adolescent Brain Cognitive Development (ABCD) Study. A psychological network approach was employed to provide a novel symptom-oriented perspective, incorporating a broad range of symptoms and brain structural variables (i.e., subcortical and cortical volume, cortical thickness, and cortical surface area). First, psychiatric symptom networks were constructed to identify the central symptoms in MDD and GAD. Second, brain structural indices were incorporated into psychiatric networks to determine the brain bridges—key brain regions linking structural anatomy to symptoms. Additionally, we identified the key intermediate symptoms that mediated the relationship between brain bridges and psychiatric networks. Given the exploratory nature of this study, we did not propose specific hypotheses.

## Methods and materials

### Participants

This study used data from a population-based youth sample collected across 21 U.S. study sites from the ongoing ABCD study (release 5.0). Each site obtained institutional review board approval before data collection. Children provided assent, and parents gave written informed consent. Data from the baseline (n = 11714, ages 9-11) and two-year follow-up (n = 10812, ages 11-14) waves were included to create a large cross-sectional sample spanning late childhood to early adolescence (ages 9-14).

High-risk adolescents were identified using the screening interview from the self-administered version of the Kiddie Schedule for Affective Disorders and Schizophrenia for DSM-5 (KSADS-5) (27). The self-administered version was developed to replicate the in-depth questioning typically conducted by a trained clinician. The screening interview has demonstrated good validity, as evidenced by strong consistency with established measures of depression and anxiety, such as the 9-item Patient Health Questionnaire and the 7-item Generalized Anxiety Disorder Scale (28). Finally, 1,568 youth were identified as at high risk for depression, and 413 were screened at high risk for anxiety (Supplementary 1). These participants then completed KSADS MDD and GAD modules to assess a range of specific symptoms and Magnetic Resonance Imaging scans to assess brain structure. Only participants who completed both brain imaging and KSADS assessments were included in the analysis.

### Symptom assessment

All current (past 2 weeks) symptoms reported by adolescents in the KSADS MDD and GAD modules were included in this study. MDD symptoms included hypersomnia, fatigue, concentration difficulties, decision-making difficulties, reduced appetite, increased appetite, agitation, slow motion, guilt, hopelessness, and self-hatred. GAD symptoms included restlessness, feeling keyed-up, fatigue, concentration difficulties, mind blanking, irritability, muscle tension, difficulty falling asleep, difficulty staying asleep, and uncontrolled worries. MDD and GAD shared overlapping symptoms in the domains of concentration difficulties, fatigue, and sleep disturbances. Participants rated the frequency of these symptoms on a scale from 0 (not at all) to 4 (nearly every day). The MDD and GAD assessments have been used in previous studies and have demonstrated good psychometric properties (28).

### MRI acquisition and processing

Details about brain imaging acquisition procedures have been detailed previously (29) (Supplementary 1). Brain data were collected on three 3T scanning systems (Siemens Prisma, Philips, and General Electric 750). Preprocessing involved correcting gradient nonlinearity distortions with scanner-specific nonlinear transformations, adjusting intensity and correcting for inhomogeneities, registering images to a custom in-house averaged reference brain in standard space, and conducting manual quality control. Cortical reconstruction and volumetric segmentation were conducted by the ABCD Data Analysis, Informatics, and Resources Center (DAIRC) using FreeSurfer v7.1.1 (https://surfer.nmr.mgh.harvard.edu/) according to standardized processing pipelines. Quality control procedures were performed by the DAIRC according to automated and manual approaches before sharing the data. Subcortical volumes were obtained using the automatic subcortical segmentation atlas in FreeSurfer, and cortical indices were obtained using the Desikan–Killiany atlas. Postprocessed subcortical and cortical volume, cortical thickness, and cortical surface area data were extracted for 50 regions of interest (ROIs). Subcortical regions included bilateral amygdala, putamen, hippocampus, thalamus, caudate, pallidum, and nuclear accumbens area. Cortical regions included bilateral precuneus, precentral cortex, lateral orbitofrontal cortex, medial orbitofrontal cortex, frontal pole, caudal middle frontal cortex, rostral middle frontal cortex, superior frontal cortex, posterior cingulate cortex, rostral anterior cingulate cortex, parahippocampal cortex, insula, inferior temporal cortex, superior temporal cortex, lingual cortex, pericalcarine cortex, postcentral cortex, and inferior parietal cortex. These ROIs were selected because prior studies in children and adolescents have shown their associations with anxiety or depression (15–23).

## Statistical analysis

### Network construction and analyses

Network construction followed the manual for psychological network analyses (30). Given the positively skewed distribution of depression and anxiety symptoms, we applied log transformation to this data prior to analysis. The effects of sex and age were controlled in the analyses. We first constructed MDD and GAD symptom networks, separately. Then, we integrated each brain structural index (i.e., subcortical and cortical volumes, cortical thickness and surface area) with MDD or GAD symptoms to create eight separate MDD-brain and GAD-brain networks.

A Gaussian graphical model was used to estimate associations between nodes, where edge weights are interpreted as partial correlation coefficients. The graphical Least Absolute Shrinkage and Selection Operator (LASSO) (31), combined with extended Bayesian information criterion (EBIC) model selection (32), was used to ensure reliable estimates. Four indices—expected influence (EI), strength, betweenness, and closeness—are commonly used to evaluate centrality in psychological networks (33). In correlation-based networks, a positive edge indicates that increased activation of one node is associated with increased activation of the connected node, whereas a negative edge indicates that increased activation of one node is associated with decreased activation of the other. EI is calculated as the sum of edge wights directly connected to a node, accounting for both positive and negative edges. It reflects the cumulative influence of a node in either activating or deactivating symptom networks. Given its ability to capture both the strength and direction of symptom interactions, EI is particularly useful in informing treatment strategies by identifying the most impactful targets for intervention (33). Strength sums the absolute values of edge weights directly connected to a node, measuring the overall connectedness. Closeness is the inverse of the average shortest path length between a node and other nodes, measuring how closely the node is linked to others. Betweenness counts the number of times that the shortest path between any two nodes passes through another node, measuring the importance of a node in linking to others.

In the symptom network, we calculated network centrality indices, which consider all network edges, to identify the central symptoms for MDD and GAD. In the MDD-brain and GAD-brain networks, we calculated bridge centrality indices, which focus exclusively on edges connecting brain and symptom nodes, to identify the brain bridges—key brain regions linking structural anatomy to symptoms (34). Additionally, we identified the strongest brain-symptom edge for each brain bridge, determining the key intermediate symptom that serves as the primary link between the brain bridges and the psychiatric networks.

To assess whether certain edges or node centralities stood out in the network, we conducted bootstrapped difference tests in an exploratory fashion to determine if their estimates differed significantly from others. The aforementioned analyses were performed using the R packages *bootnet* (version 1.6) (30) and *qgraph* (version 1.9.8) (35) in R version 4.4.0. Layouts for each graph were determined by the Fruchterman-Reingold algorithm (36).

### Estimation of edge accuracy and network stability

We first assessed the accuracy of edge weights by calculating their 95% confidence intervals (CIs) derived from 1000 non-parametric bootstrap samples. Second, we evaluated the stability of centrality indices using a case-dropping subset bootstrap procedure with 1000 samplings. Centrality indices were repeatedly calculated from subsets of data with an increasing proportion of subjects removed. Stability was quantified by calculating the correlation stability coefficient, representing the maximum proportion of cases that can be removed from the sample while retaining, with 95% certainty. The correlation between the original centrality indices (full sample) and those derived from subsamples needs to be at least 0.7. The estimations were conducted using the R package *bootnet* (version 1.6).

## Results

A total of 1,568 adolescents at risk for MDD and 413 adolescents at risk for GAD were included in the analyses. The results were organized according to the reporting standards for psychological networks (37). Demographic and clinical characteristics of the risk samples are provided in Table 1.

**Table 1.**
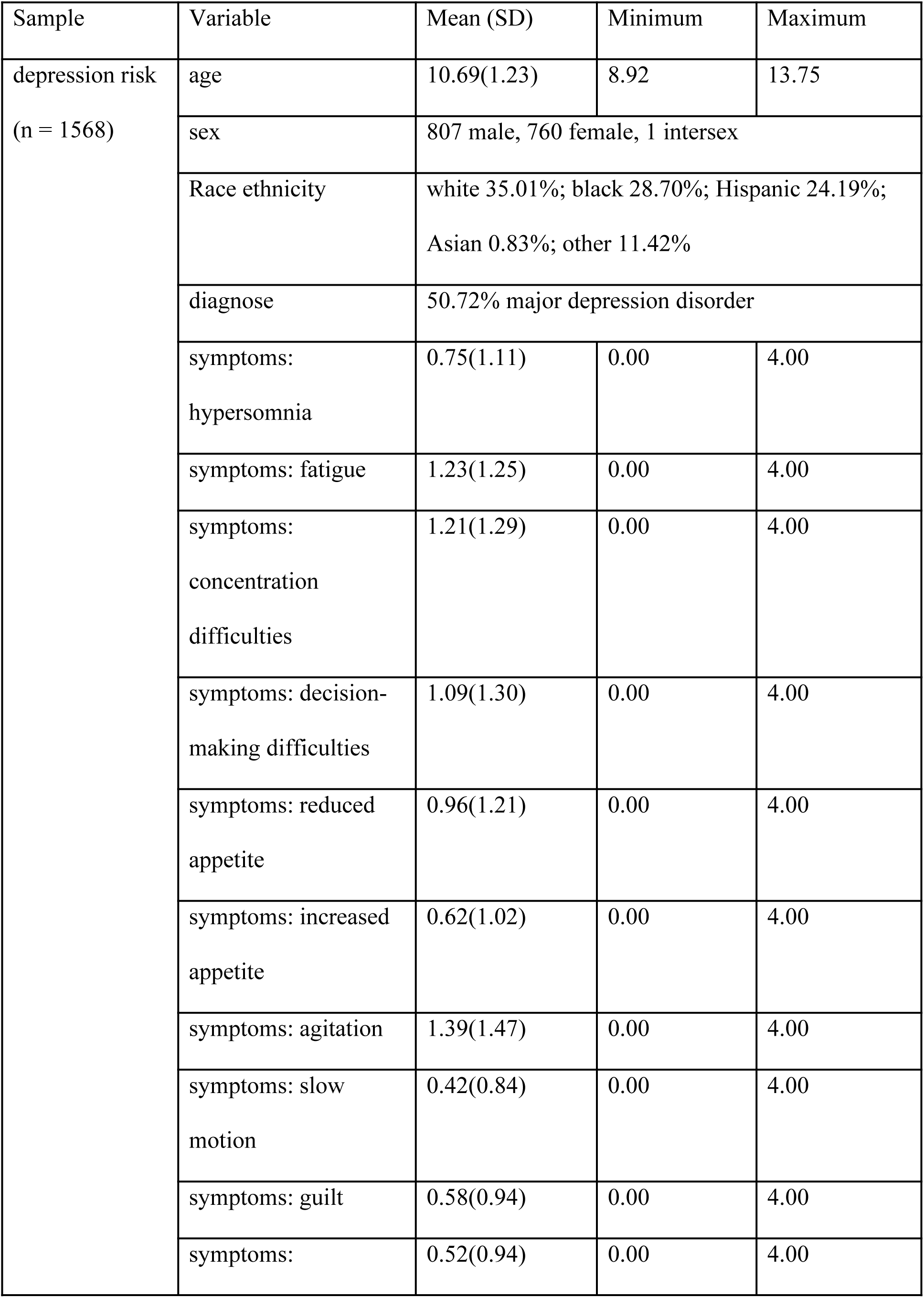
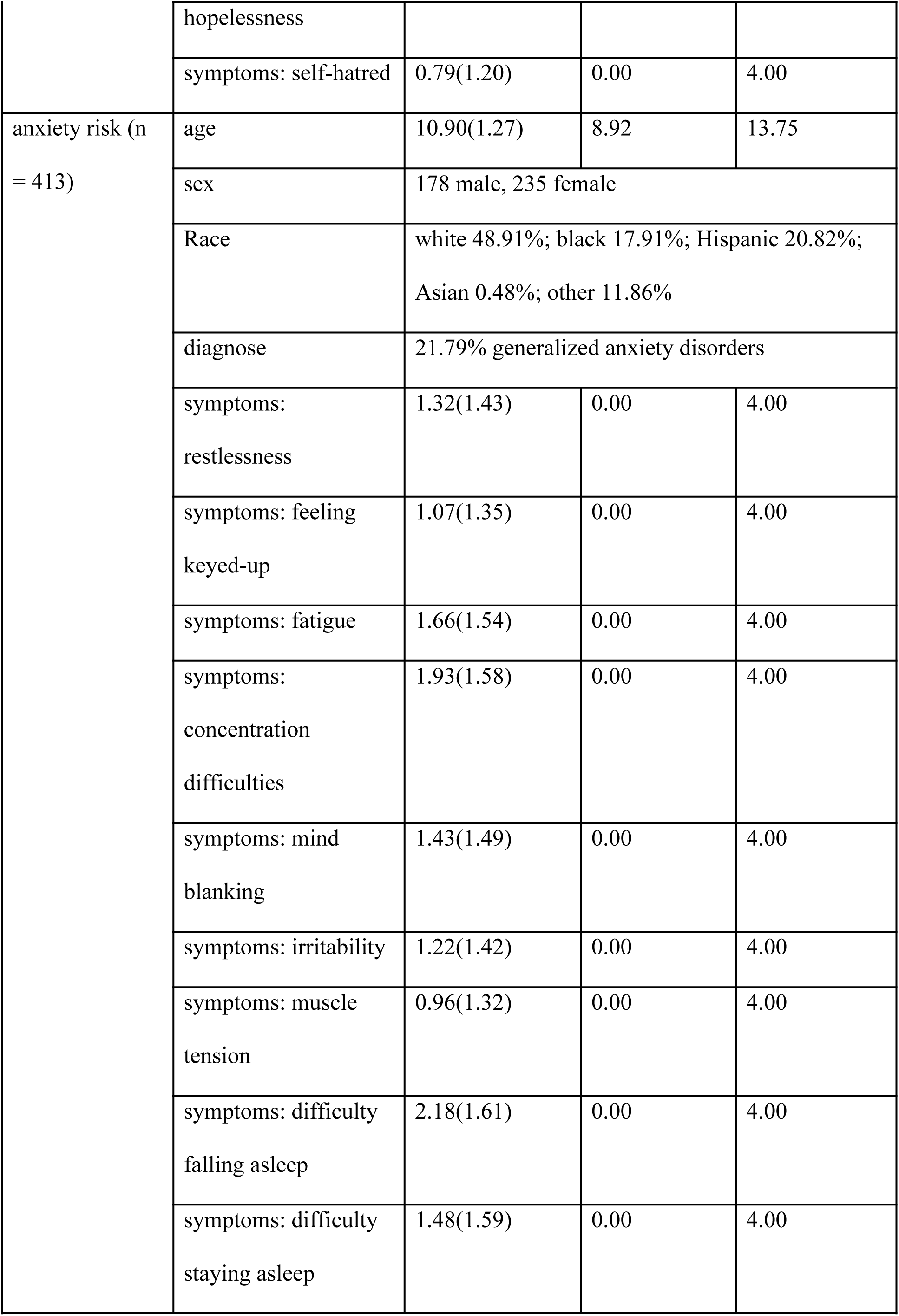
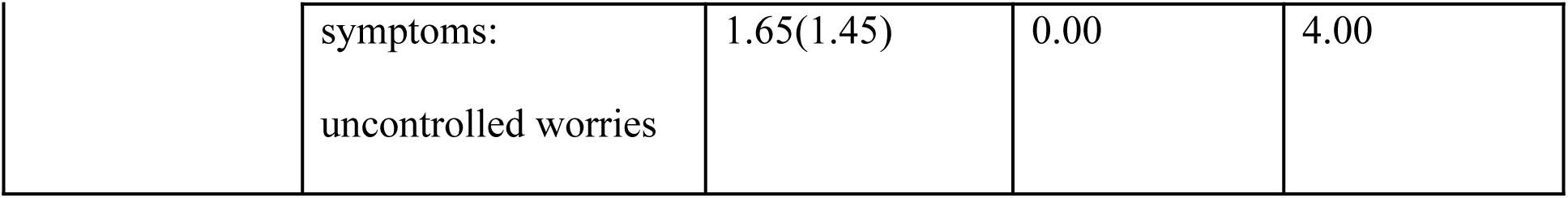
Demographic and clinical characteristics.

### Network stability checks

Sample edge weights were comparable to bootstrap mean values (Supplementary 1 Figure S1), indicating that edges were estimated accurately. In the symptom-only networks, the stability coefficients of centrality indices—except for betweenness—were above 0.25, indicating high stability. In the symptom-brain networks, EI demonstrated higher stability than other indices. Given its relevance in clinical practices (see Methods and materials), EI was selected as the primary centrality measure for identifying brain bridges. EI stability in the MDD–brain networks was acceptable across all structural indices except cortical thickness. However, all EI stability for the GAD–brain networks fell below the 0.25 threshold, indicating limited stability. A lower stability coefficient suggests that the interpretations of centrality rankings based on that index should be approached with caution. To address potential risks in the interpretation of rankings, we supplemented the results from bootstrapped difference tests for brain edges and EI in all the networks to strengthen confidence in our findings (37).

### Central symptoms in MDD and GAD symptom networks

Figure 1 presents the MDD and GAD symptom networks along with their centrality. Symptoms in these networks were generally positively connected. The strongest edge was *hopelessness*–*self-hatred* (*r* = 0.340) in the MDD network and *difficulty falling asleep*–*difficulty staying asleep* (*r* = 0.317) in the GAD network (Supplementary 2 Tables S1-S2). These edges stood out as significantly different from most other edges (Supplementary 1 Figure S2).

**Figure 1.**
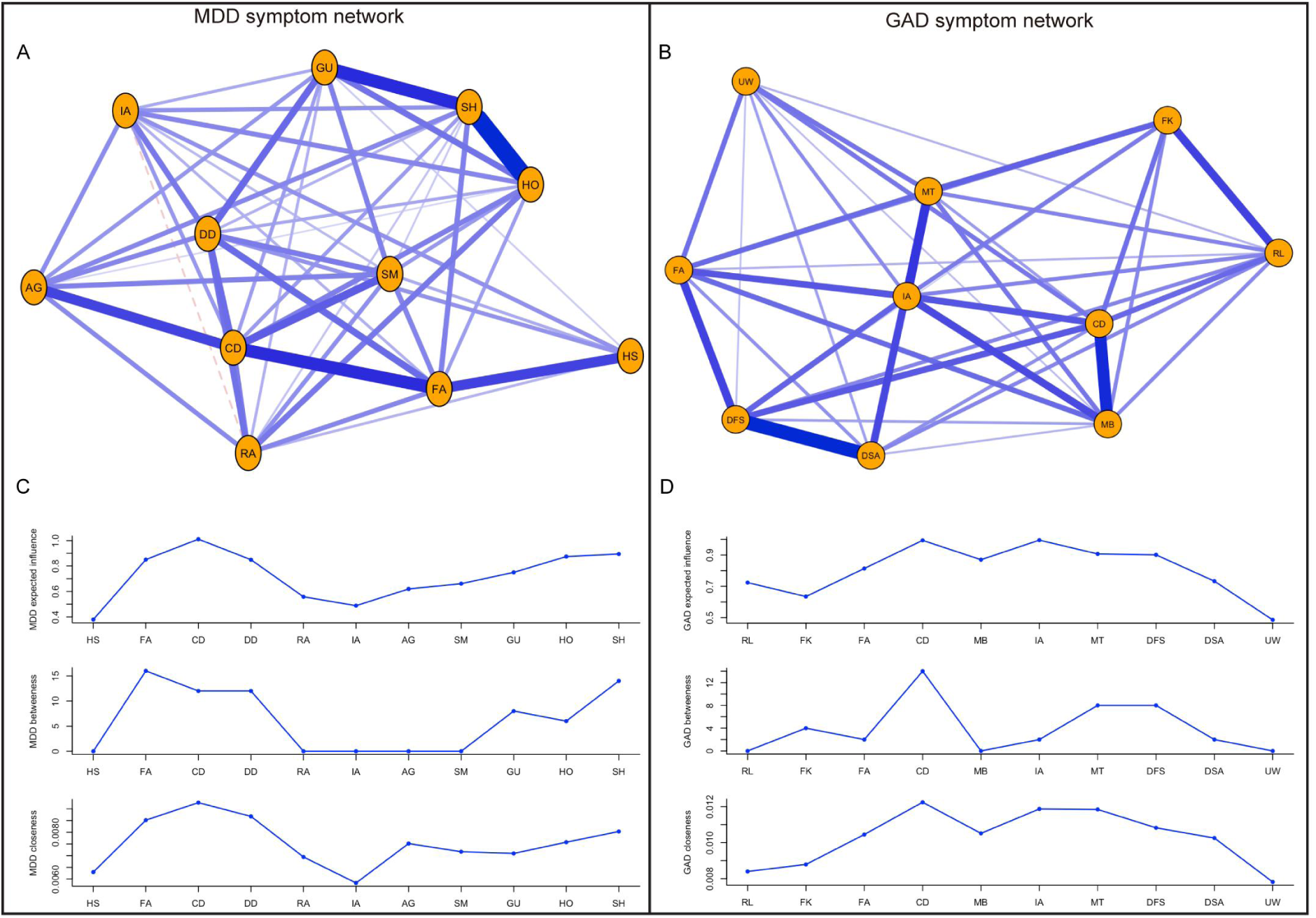
MDD and GAD symptom networks and network centrality. No cutoff value was applied to network visualizations. Abbreviations: HS, hypersomnia; FA, fatigue; CD, concentration difficulties; DD, decision-making difficulties; RA, reduced appetite; IA, increased appetite; AG, agitation; SM, slow motion; GU, guilt; HO, hopelessness; SH, self-hatred; RL, restlessness; FK, feeling keyed-up; MB, mind blanking; IA, irritability; MT, muscle tension; DFS, difficulty falling asleep; DSA, difficulty staying asleep; UW, uncontrolled worries.

Regarding centrality, *concentration difficulties* showed the highest influence in the MDD network (EI = 1.005), significantly different from most other symptoms (Supplementary 1 Figure S2). It was also the symptom most closely related to other symptoms (closeness = 0.009). *Self-hatred* also demonstrated higher centrality (EI = 0.926). For GAD, the most influential symptoms were *irritability* (EI = 0.984) and *concentration difficulties* (EI = 0.980), both significantly different from most other symptoms (Supplementary 1 Figure S2). *Concentration difficulties* was also the symptom most closely related to other symptoms (closeness = 0.012). All centrality estimates are shown in Tables S3-S4 in Supplementary 2.

### Brain bridges and their intermediate symptoms in MDD-Brain and GAD-Brain networks

Intra-relations within brain and symptom nodes were closer than inter-relations between brain and symptom nodes (Supplementary 1 Figure S3; Supplementary 2 Tables S5-S12). The MDD- and GAD-brain networks and bridge EI regarding subcortical structures are presented in Figure 2, with only brain-symptom edges and brain bridge centrality displayed for clarity. The MDD-brain network exhibited more significant edges than the GAD-brain network, indicating stronger associations with subcortical structures. The brain-symptom edge weights ranged from –0.040 to 0.048 for MDD (53.85% negative edges) and ranged from –0.032 to 0.046 for GAD (30.00% negative edges; Supplementary 2 Tables S5-S6). The *left-pallidum* (EI = –0.065) and *right-pallidum* volumes (EI = 0.077) showed the highest bridge centrality in connection with MDD symptoms (Supplementary 2 Table S13), significantly different from many other nodes (Supplementary 1 Figure S4). Notably, the *left-pallidum* was negatively associated with *self-hatred* (*r* = –0.040), while the *right-pallidum* was positively associated with *hypersomnia* (*r* = 0.048). The *left-nucleus accumbens (NAcc)* volume (EI = 0.102) exhibited the highest bridge centrality in connection with GAD symptoms (Supplementary 2 Table S14), significantly different from most other nodes (Supplementary 1 Figure S5). Specifically, the *left-NAcc* volume was mainly connected to *restlessness* (*r* = 0.046).

**Figure 2.**
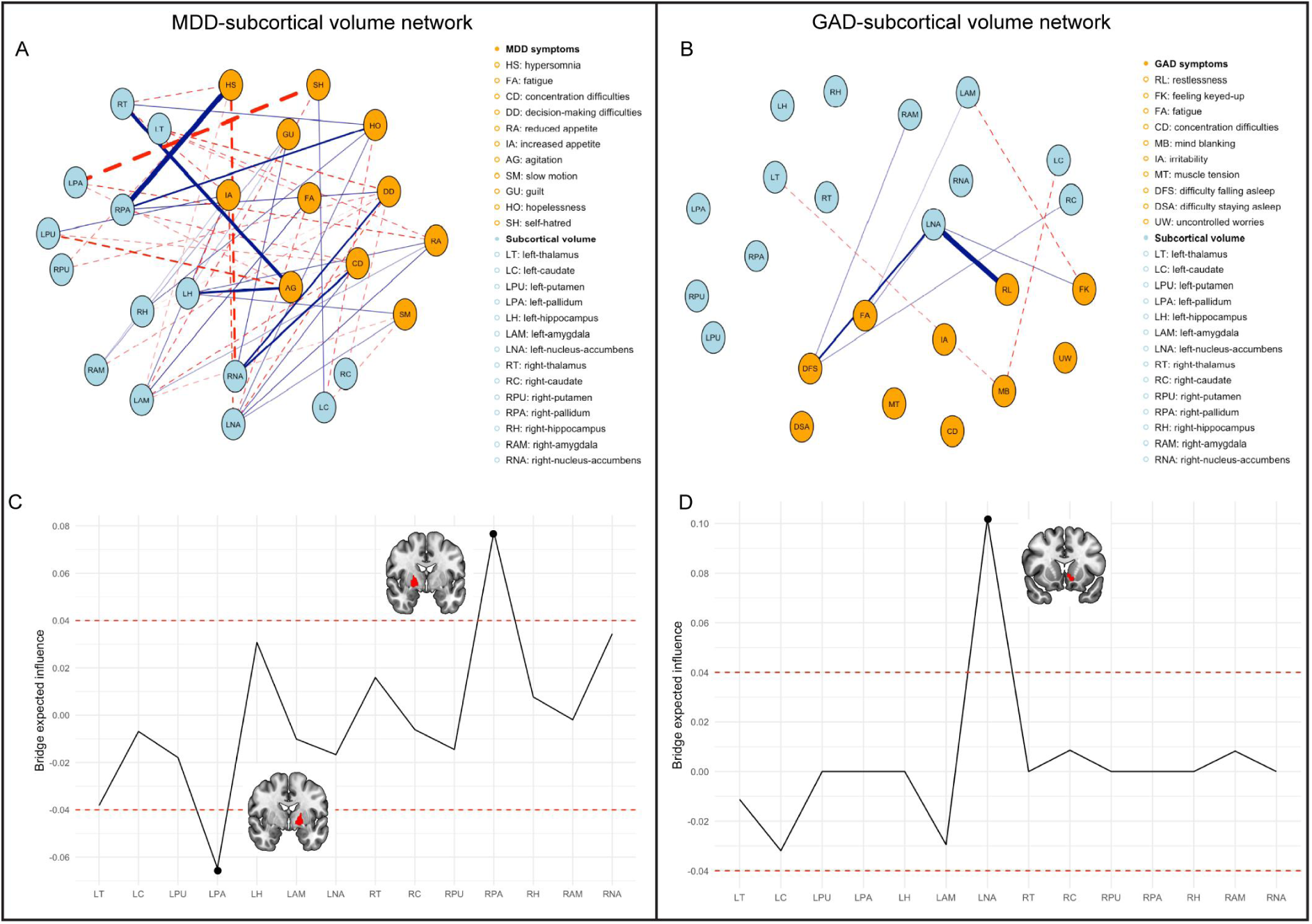
Brain-symptom networks and brain bridge centrality for subcortical volume. Only brain-symptom edges were presented, with no specific cutoff values applied for network visualization. The red dashed lines in bridge centrality plot represent values of 0.04 and –0.04 for clarity.

Regarding cortical structures, the brain-symptom edges were relatively sparse and weak (Supplementary 2 Tables S7-S12). Cortical volume, thickness, and surface area were considered collectively to describe the most influential cortical structures associated with MDD and GAD symptoms, while specific intermediate symptoms were not determined. Figure 3 shows the bridge EI of cortical structures, with all values provided in Tables S15-S20 in Supplementary 2. The bridge cortical structures connecting to MDD and GAD symptoms mostly converge on the right frontal pole. However, its EI was not significantly different from other nodes in the network and should be interpreted with caution (Supplementary 1 Figures S4-S5).

**Figure 3.**
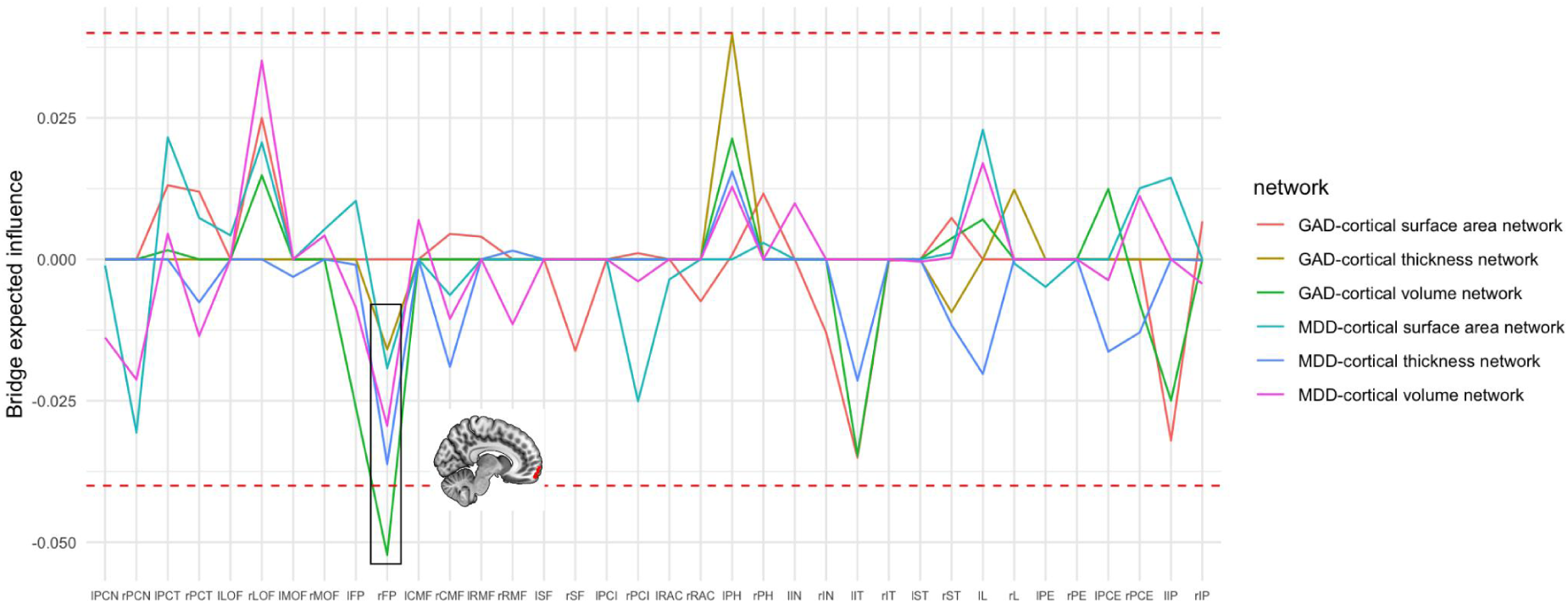
Brain bridge centrality for cortical structures. The red dashed lines represent EI values of 0.04 and –0.04 for clarity. Abbreviations: lPCN, left-precuneus; rPCN, right-precuneus; lPCT, left-precentral cortex; rPCT, right-precentral cortex; lLOF, left-lateral orbitofrontal cortex; rLOF, right-lateral orbitofrontal cortex; lMOF, left-medial orbitofrontal cortex; rMOF, right-medial orbitofrontal cortex; lFP, left-frontal pole; rFP, right-frontal pole; lCMF, left-caudal middle frontal cortex; rCMF, right-caudal middle frontal cortex; lRMF, left-rostral middle frontal cortex; rRMF, right-rostral middle frontal cortex; lSF, left-superior frontal cortex; rSF, right-superior frontal cortex, lPCI, left-posterior cingulate cortex; rPCI, right-posterior cingulate cortex; lRAC, left-rostral anterior cingulate cortex; rRAC, right-rostral anterior cingulate cortex; lPH, left-parahippocampal cortex; rPH, right-parahippocampal cortex; lIN, left-insula; rIN, right-insula; lIT, left-inferior temporal cortex; rIT, right-inferior temporal cortex; lST, left-superior temporal cortex; rST, right-superior temporal cortex; lL, left-lingual cortex; rL, right-lingual cortex; lPE, left-pericalcarine cortex; rPE, right-pericalcarine cortex, lPCE, left-postcentral cortex; rPCE, right-postcentral cortex; lIP, left-inferior parietal cortex, rIP, right-inferior parietal cortex.

## Discussion

This study examined the symptom structure of MDD and GAD and their symptom-oriented neural substrates in adolescents at risk. First, our results indicated that concentration difficulties emerged as a shared core symptom across both disorders, while self-hatred and irritability were uniquely central to MDD and GAD, respectively. Second, we identified that pallidum volume was the most influential subcortical structure in MDD symptoms, whereas left-NAcc volume had the strongest influence in GAD symptoms. These brain bridges exerted their influence on the psychiatric network through distinct symptom pathways. By analyzing individual symptoms rather than aggregated measures, these findings provide novel insights into the neurobiological underpinnings of MDD and GAD.

In high-risk youth, concentration difficulties emerged as the core symptom for both MDD and GAD, a finding that diverges from studies in the general community, where it is often considered peripheral (10–14,38–39). This is particularly significant given that children and adolescents are at a critical stage in their cognitive development, and concentration is crucial for academic success and social interactions (40,41). Impaired concentration during this period may cause greater distress, leading to higher symptom severity. Notably, among the overlapping symptoms (i.e., concentration difficulties, fatigue, and sleep disturbances), concentration difficulties emerged as a shared core symptom for MDD and GAD. It may reflect its unique role in underlying a common latent cognitive vulnerability that contributes to their co-occurrence. Additionally, self-hatred emerged as a central symptom in the MDD network, while irritability was central in the GAD network. These findings emphasize the differential characteristics of each disorder, consistent with previous studies highlighting the central role of self-hatred in depression (13,14,38,39) and of irritability in anxiety (10).

Among subcortical structures, pallidum volume played the most influential role in connecting to MDD symptoms. The pallidum is known to be involved in reward and motivation behavior (42,43), both of which are key features of youth depression (44). A previous study reported that higher depressive symptoms were associated with larger total pallidal volume in children and adolescents (19); however, it did not distinguish between hemispheres. When examined separately, we observed a significant lateralized effect: smaller left-pallidum volume and larger right-pallidum volume were each associated with higher symptom levels. In the brain-symptom network, smaller left-pallidum volume was primarily associated with increased symptom of self-hatred. This finding aligns with previous work linking neural activity in the left-pallidum to self-esteem (45). In contrast, larger right-pallidum volume was associated with the symptom of hypersomnia. A finding from the UK Biobank also linked a larger right-pallidum volume with longer sleep duration (46). These lateralized effects suggest distinct pathways through which the pallidum may contribute to depression symptomatology, warranting further investigation into its role beyond reward processing and motivation.

Left-NAcc volume emerged as the most influential subcortical structure in the GAD network. The NAcc is a critical component of the brain’s reward system and plays a key role in suppressing irrelevant or nonrewarded actions to facilitate goal-directed behavior (47). Previous studies in adolescents have reported mixed findings regarding the association between NAcc volume and anxiety (17–20). This inconsistency may stem from the complex interactions among individual symptoms and brain regions, which aggregated symptom scores may fail to capture. In the present study, we extended prior work by identifying the cumulative influence of left-NAcc volume on GAD symptoms. Importantly, we found that NAcc volume primarily influenced the GAD network through the symptom of restlessness, which suggests that alterations in the NAcc may impair the ability to suppress irrelevant motor actions, leading to heightened physical agitation. These findings align with research indicating that somatic anxiety is associated with reduced goal-directed exploration (48), further supporting the NAcc’s role in regulating motor and cognitive responses to anxiety-provoking situations (49).

Regarding cortical structures, MDD and GAD symptoms were primarily negatively associated with the right-frontal pole, a region encompassing the ventromedial prefrontal cortex (vmPFC). The vmPFC is crucial for various social, cognitive, and affective functions, many of which are disrupted in depression and anxiety (50). By considering a comprehensive range of symptoms and their interactions, our research helps clarify the conflicting findings regarding the association between adolescent anxiety and depression and vmPFC volume or thickness (15, 16). Our findings also suggest that structural abnormalities in the right-frontal pole may be a potential neural marker underlying the comorbidity of MDD and GAD. However, as the brain-symptom networks regarding cortical structures were sparse and weak in our study, this interpretation requires further study for substantiation.

### Implications and limitations

By adopting a network perspective to map the dynamic interactions between symptoms and brain anatomy, this study provides significant clinical implications. For instance, targeting core symptoms—concentration difficulties and self-hatred in MDD, and concentration difficulties and irritability in GAD—may facilitate an overall deactivation of subclinical symptom networks, presenting a promising approach for early intervention. Notably, addressing concentration difficulties could be particularly effective in preventing the comorbidity of MDD and GAD. Regarding brain bridges, the pallidum and left-NAcc emerged as key neural connectors between brain structures and symptom networks, underscoring the critical role of reward circuitry development in shaping depression and anxiety. These regions may serve as potential biomarkers for the early identification of individuals at risk. From a network perspective, intervening in these brain bridges could contribute to a global reduction in psychiatric network activation. Brain-based treatments such as neuromodulation may prioritize these regions as treatment targets. For example, deep brain stimulation targeting the reward circuitry has been used in previous research to treat individual with depression and anxiety disorders (51–54).

This study is the first to examine the neural underpinnings of MDD and GAD from a symptom-oriented perspective. However, several limitations should be acknowledged. First, although the adolescents in our study (identified from over 10,000 community adolescents) represent a relatively large high-risk group, the sample size may still be insufficient for reliably estimating GAD-brain networks involving cortical structures. To combat this issue effectively, we have employed the LASSO regularization procedure with EBIC selection to ensure robust estimates, a method especially recommended for relatively small psychological datasets (30), and bootstrap difference tests to strengthen our results. Nevertheless, larger samples are still needed to replicate these findings. Second, although centrality index of EI provides an overall measure of a brain node’s influence on symptom networks, the cross-sectional design of this study limits our ability to draw causal conclusions. In the ABCD Study Release 5.0, KSADS assessments are available only for the baseline and the two-year follow-up waves. In our study, data from both waves were combined to ensure a sufficient sample size of at-risk youth for analysis. As additional longitudinal data become available, future research will be better positioned to investigate causal relationships between individual symptoms and atypical brain development. Third, our focus is on identifying biomarkers within a risk sample, where comorbidity between disorders—or with other conditions—is naturally present. To better approach a real-world context, we did not exclude participants with diagnosed or comorbid conditions. Nevertheless, it will also be important for future research to examine biomarkers within clinically diagnosed groups, specifically for individuals with GAD, MDD, and their comorbidity. Finally, the brain-symptom links identified in this study may reflect the shared genetic and environmental influences shaping both psychiatric symptoms and brain development. The underlying mechanisms driving these associations warrant further investigation.

### Conclusion

This study investigated the network structure of MDD and GAD symptoms and identified key brain structures underlying these symptoms in risk adolescents. Concentration difficulties emerged as a shared core symptom across both disorders, while self-hatred and irritability were uniquely central to MDD and GAD, respectively. Notably, we identified the brain bridge role of left- and right-pallidum volume in MDD and left-NAcc volume in GAD. These findings provide novel insights into early symptom-oriented biomarkers and have important implications for designing targeted preventive interventions.

## Author contributions

QD, SW, and DR designed the study. QD and SP carried out the statistical analysis and produced figures. All authors interpreted the results. QD wrote the first draft of the manuscript and all authors contributed to editing and commenting on the final version. SW supervised the study.

## Data availability statement

Only researchers with an approved NDA Data Use Certification (DUC) may obtain ABCD Study data. ABCD Study data can be obtained by applying to the NDA directly here: https://nda.nih.gov/abcd/

