## Supplementary 1 for "Neuroanatomical signatures of depression and anxiety in at-risk adolescents: A symptom-oriented perspective"

**High risk sample**

For the MDD high-risk sample, the baseline wave included 1001 adolescents, and the two-year follow-up wave included 715 adolescents. One hundred and forty-eight adolescents participated in both waves, for which only baseline data were included in the analysis. For the GAD high-risk sample, the baseline wave included 225 adolescents, and the two-year follow-up wave included 209 adolescents. Twenty-one adolescents participated in both waves, for which only baseline data were included in the analysis.

**MRI acquisition and processing**

The T1 weighted images were acquired using a three-dimensional magnetization-prepared rapid-acquisition gradient echo sequence with a voxel size of 1 mm3 and a T2 weighted axial fast-spoiled gradient echo sequence (repetition time [TR] = 2400–2500 milliseconds, echo time [TE] = 2–2.9 milliseconds, matrix size = 256 × 256, field of view = 256 × 240–256 mm2, flip angle = 8°, inversion delay = 1060 milliseconds, and 176–225 sections).

Table S1 Centrality stability for the networks.

| **Networks** | **Network EI** | **Network strength** | **Network closeness** | **Network betweenness** |
| --- | --- | --- | --- | --- |
| MDD symptom network | 0.750 | 0.750 | 0.594 | 0.205 |
| GAD symptom network | 0.596 | 0.516 | 0.361 | 0 |
|  | **Bridge EI** | **Bridge strength** | **Bridge closeness** | **Bridge betweenness** |
| MDD-subcortical volume network | 0.750 | 0.750 | 0.673 | 0 |
| MDD-cortical volume network | 0.284 | 0.205 | 0 | 0 |
| MDD-cortical thickness network | 0.205 | 0.128 | 0.050 | 0.050 |
| MDD-cortical surface area network | 0.361 | 0.284 | 0.050 | 0.050 |
| GAD-subcortical volume network | 0.206 | 0.128 | 0 | 0.049 |
| GAD-cortical volume network | 0.128 | 0.049 | 0 | 0 |
| GAD-cortical thickness network | 0.049 | 0.049 | 0.049 | 0.049 |
| GAD-cortical surface area network | 0.049 | 0 | 0 | 0 |

Figure S1 Bootstrapped 95%CI for edge weights.

Abbreviations: HS, hypersomnia; FA, fatigue; CD, concentration difficulties; DD, decision-making difficulties; RA, reduced appetite; IA, increased appetite; AG, agitation; SM, slow motion; GU, guilt; HO, hopelessness; SH, self-hatred; RL, restlessness; FK, feeling keyed-up; MB, mind blanking; IA, irritability; MT, muscle tension; DFS, difficulty falling asleep; DSA, difficulty staying asleep; UW, uncontrolled worries; LT, left-thalamus; LC, left-caudate; LPU, left-putamen; LPA, left-pallidum; LH, left-hippocampus; LAM, left-amygdala; LNA, left-nucleus-accumbens; RT, right-thalamus; RC, right-caudate; RPU, right-putamen; RPA, right-pallidum; RH, right-hippocampus; RAM, right-amygdala; RNA, right-nucleus-accumbens; lPCN, left-precuneus; rPCN, right-precuneus; lPCT, left-precentral cortex; rPCT, right-precentral cortex; lLOF, left-lateral orbitofrontal cortex; rLOF, right-lateral orbitofrontal cortex; lMOF, left-medial orbitofrontal cortex; rMOF, right-medial orbitofrontal cortex; lFP, left-frontal pole; rFP, right-frontal pole; lCMF, left-caudal middle frontal cortex; rCMF, right-caudal middle frontal cortex; lRMF, left-rostral middle frontal cortex; rRMF, right-rostral middle frontal cortex; lSF, left-superior frontal cortex; rSF, right-superior frontal cortex, lPCI, left-posterior cingulate cortex; rPCI, right-posterior cingulate cortex; lRAC, left-rostral anterior cingulate cortex; rRAC, right-rostral anterior cingulate cortex; lPH, left-parahippocampal cortex; rPH, right-parahippocampal cortex; lIN, left-insula; rIN, right-insula; lIT, left-inferior temporal cortex; rIT, right-inferior temporal cortex; lST, left-superior temporal cortex; rST, right-superior temporal cortex; lL, left-lingual cortex; rL, right-lingual cortex; lPE, left-pericalcarine cortex; rPE, right-pericalcarine cortex, lPCE, left-postcentral cortex; rPCE, right-postcentral cortex; lIP, left-inferior parietal cortex, rIP, right-inferior parietal cortex.

Figure S2. Bootstrapped difference tests for edge-weights and network expected influences in MDD and GAD symptom networks.

Abbreviations: HS, hypersomnia; FA, fatigue; CD, concentration difficulties; DD, decision-making difficulties; RA, reduced appetite; IA, increased appetite; AG, agitation; SM, slow motion; GU, guilt; HO, hopelessness; SH, self-hatred; RL, restlessness; FK, feeling keyed-up; MB, mind blanking; IA, irritability; MT, muscle tension; DFS, difficulty falling asleep; DSA, difficulty staying asleep; UW, uncontrolled worries.

Figure S3 MDD-brain networks and GAD-brain networks with all edges.

Figure S4. Bootstrapped difference tests for edge-weights and bridge expected influences in MDD-brain networks.

Abbreviations: HS, hypersomnia; FA, fatigue; CD, concentration difficulties; DD, decision-making difficulties; RA, reduced appetite; IA, increased appetite; AG, agitation; SM, slow motion; GU, guilt; HO, hopelessness; SH, self-hatred; LT, left-thalamus; LC, left-caudate; LPU, left-putamen; LPA, left-pallidum; LH, left-hippocampus; LAM, left-amygdala; LNA, left-nucleus-accumbens; RT, right-thalamus; RC, right-caudate; RPU, right-putamen; RPA, right-pallidum; RH, right-hippocampus; RAM, right-amygdala; RNA, right-nucleus-accumbens; lPCN, left-precuneus; rPCN, right-precuneus; lPCT, left-precentral cortex; rPCT, right-precentral cortex; lLOF, left-lateral orbitofrontal cortex; rLOF, right-lateral orbitofrontal cortex; lMOF, left-medial orbitofrontal cortex; rMOF, right-medial orbitofrontal cortex; lFP, left-frontal pole; rFP, right-frontal pole; lCMF, left-caudal middle frontal cortex; rCMF, right-caudal middle frontal cortex; lRMF, left-rostral middle frontal cortex; rRMF, right-rostral middle frontal cortex; lSF, left-superior frontal cortex; rSF, right-superior frontal cortex, lPCI, left-posterior cingulate cortex; rPCI, right-posterior cingulate cortex; lRAC, left-rostral anterior cingulate cortex; rRAC, right-rostral anterior cingulate cortex; lPH, left-parahippocampal cortex; rPH, right-parahippocampal cortex; lIN, left-insula; rIN, right-insula; lIT, left-inferior temporal cortex; rIT, right-inferior temporal cortex; lST, left-superior temporal cortex; rST, right-superior temporal cortex; lL, left-lingual cortex; rL, right-lingual cortex; lPE, left-pericalcarine cortex; rPE, right-pericalcarine cortex, lPCE, left-postcentral cortex; rPCE, right-postcentral cortex; lIP, left-inferior parietal cortex, rIP, right-inferior parietal cortex.

Figure S5 Bootstrapped difference tests for edge-weights and bridge expected influences in GAD-brain networks.

Abbreviations: RL, restlessness; FK, feeling keyed-up; FA, fatigue; CD, concentration difficulties; MB, mind blanking; IA. irritability; MT, muscle tension; DFS, difficulty falling asleep; DSA, difficulty staying asleep; UW, uncontrolled worries; LT, left-thalamus; LC, left-caudate; LPU, left-putamen; LPA, left-pallidum; LH, left-hippocampus; LAM, left-amygdala; LNA, left-nucleus-accumbens; RT, right-thalamus; RC, right-caudate; RPU, right-putamen; RPA, right-pallidum; RH, right-hippocampus; RAM, right-amygdala; RNA, right-nucleus-accumbens; lPCN, left-precuneus; rPCN, right-precuneus; lPCT, left-precentral cortex; rPCT, right-precentral cortex; lLOF, left-lateral orbitofrontal cortex; rLOF, right-lateral orbitofrontal cortex; lMOF, left-medial orbitofrontal cortex; rMOF, right-medial orbitofrontal cortex; lFP, left-frontal pole; rFP, right-frontal pole; lCMF, left-caudal middle frontal cortex; rCMF, right-caudal middle frontal cortex; lRMF, left-rostral middle frontal cortex; rRMF, right-rostral middle frontal cortex; lSF, left-superior frontal cortex; rSF, right-superior frontal cortex, lPCI, left-posterior cingulate cortex; rPCI, right-posterior cingulate cortex; lRAC, left-rostral anterior cingulate cortex; rRAC, right-rostral anterior cingulate cortex; lPH, left-parahippocampal cortex; rPH, right-parahippocampal cortex; lIN, left-insula; rIN, right-insula; lIT, left-inferior temporal cortex; rIT, right-inferior temporal cortex; lST, left-superior temporal cortex; rST, right-superior temporal cortex; lL, left-lingual cortex; rL, right-lingual cortex; lPE, left-pericalcarine cortex; rPE, right-pericalcarine cortex, lPCE, left-postcentral cortex; rPCE, right-postcentral cortex; lIP, left-inferior parietal cortex, rIP, right-inferior parietal cortex.
